# Kinetics of i-motif folding within the duplex context

**DOI:** 10.1101/2024.08.18.608433

**Authors:** Rugiya Alieva, Anna Keshek, Timofei Zatsepin, Victor Orlov, Andrey Aralov, Elena Zavyalova

**Affiliations:** Chemistry Department of Lomonosov Moscow State University, Moscow, Russia; Belozersky Institute of Physico-Chemical Biology of Lomonosov Moscow State University, Moscow, Russia; Shemyakin-Ovchinnikov Institute of Bioorganic Chemistry, Russian Academy of Sciences, Moscow, Russia; RUDN University, Moscow, Russia

**Keywords:** DNA, i-motif, pH dependence, stopped-flow kinetics

## Abstract

Non-canonical nucleic acid structures possess an ability to interact selectively with proteins, thereby exerting influence over various intracellular processes. Numerous studies indicate that genomic G-quadruplexes and i-motifs are involved in the regulation of transcription. These structures are formed temporarily during the unwinding of the DNA double helix; and their direct determination is a rather difficult task. In addition, i-motif folding is pH-dependent, with most i-motifs having low stability at neutral pH. However, some genomic i-motifs with long cytosine repeats were shown to be stable at pH 7.3, suggesting their functionality within the nucleus. Here we studied pH-dependent behavior of a model i-motif with flanking sequences that forms a duplex motif. Kinetic studies on bimodular structures with cytosine residues replaced with an environment-sensitive fluorescent label reveal the stabilization of the i-motif structure near the i-motif-duplex junction. These results highlight the importance of the natural environment of i-motifs for the correct assessment of their stability.

## Introduction

Non-canonical nucleic acid structures provide a variety of effects at both DNA and RNA levels. Several comprehensive reviews summarize known examples of the regulatory role of these structures as well as associated genetic disorders [1-3]. i-Motifs are among the significant non-canonical nucleic acid structures, but are much less studied compared to triplexes or G-quadruplexes. Their structure is formed by two parallel duplexes intercalated in the antiparallel orientation and held together by hemi-protonated cytosine-cytosine^+^ (C:C^+^) base pairs (Figure 1). i-Motifs are formed by the oligonucleotides bearing cytosine repeats at slightly acidic pH. Typically, transition pH values (pH_i_, the pH value at which 50% of the DNA is folded into an i-motif) are in the 5 – 6 range [4-6]. iMs with extended cytosine repeats are stable even at neutral pH having pH_i_ as high as 7.9 [7,8]. However, folding of the i-motif in the presence of a complementary strand was not observed even for relatively stable i-motif sequences; additional stressing conditions are required to destabilize the DNA duplex [9,10]. A recent study has shown that i-motif formation concerns only a tiny fraction (possibly 1%) of genomic sites with i-motif formation propensity [11].

**Figure 1.**
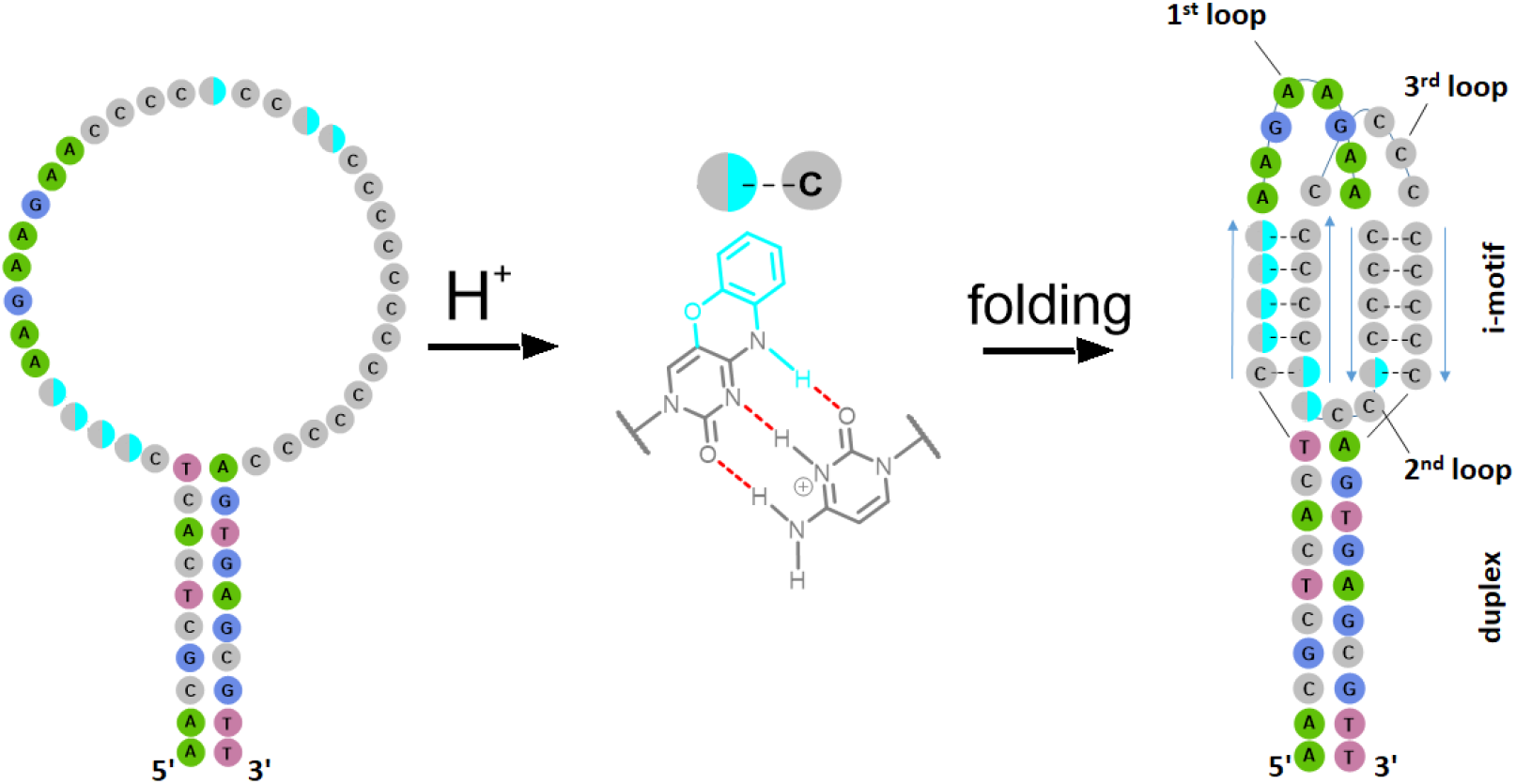
pH-dependent structure of aptamer BV42. Two cytosines form a pair coordinating one proton. Several hemiprotonated cytosine pairs form the i-motif structure with a geometry of two intercalated parallel duplexes. The structure of a cytosine mimic, 1,3-diaza-2-oxophenoxazine, is shown in comparison with cytosine. A single modification was introduced in one of the cyan-labeled sites.

Regulatory proteins provide another way to regulate the i-motif folding allowing their functionality. For example, an i-motif binding regulatory protein (BmILF) was identified. This protein specifically bound to the i-motif and activated transcription of the *BmPOUM2* gene [12]. On the contrary, regulatory protein hnRNP binds to the i-motif structure in the *BCL2* promoter region and unfolds it, increasing transcription efficiency [13]. Both examples promote further elucidation of the peculiarities of i-motif folding in a duplex context. Here we studied an artificial i-motif-duplex structure in order to investigate the influence of the duplex part on i-motif folding kinetics. Similar studies were reported previously for G-quadruplexes in the duplex context [14-16], whereas the i-motif-duplex context remains under-researched.

Based on an *in-silico* approach, BV42, a DNA aptamer to hemagglutinin of influenza A virus, was constructed [17]. BV42 is a bimodular structure where cytosine-rich stretch can fold into the i-motif at neutral conditions with pH_i_ of 7.1 (Figure 1) [18]. The i-motif structure of BV42 is responsible for hemagglutinin binding; an unfolded i-motif is unable to recognize the protein. Thus, BV42 is a switchable DNA-based recognizing element with pH-dependent behavior. The i-motif structure is flanked by complementary strands that form duplex module with an i-motif-duplex interface. Interestingly, the terminal pair in the duplex module near the i-motif-duplex junction is unassembled, resulting in the formation of a flexible hinge structure [19]. The hinge structure determines the formation of multiple conformers with different angles between the axes of the two modules. Here, the kinetics of i-motif folding and unfolding were studied to give insight into the stages of i-motif assembly in the duplex context.

## Results and Discussion

### Folding and unfolding of the i-motif during pH changes

The cytosine mimic, 1,3-diaza-2-oxophenoxazine (tC°, Figure 2) preserves the cytosine H-bonding pattern and minimally affects iM structure and stability [19]. Moreover, its microenvironment-dependent fluorescent properties allow for the kinetic study of i-motif assembly and monitoring of i-motif folding in cells [20-22]. Several cytosines were chosen for modification in different parts of the i-motif; a set of aptamers with a single modification (Table 1) was synthesized.

**Table 1.** Sequences and sites of modification within the DNA aptamers studied in this work. **X** is 1,3-diaza-2-oxophenoxazine modification (tC°). Complementary sequences are underlined. Loops are shown in bold italics.

| Code | Sequence (5'→3') |
| --- | --- |
| BV42 | <u>AACGCTCACT</u> CCCCC **AAGAAGAA** CCCCC **CCC** CCCCC **CCCC** CCCCC<br><u>AGTGAGCGTT</u> |
| BV42°C12 | <u>AACGCTCACT</u> CXCCC **AAGAAGAA** CCCCC **CCC** CCCCC **CCCC** CCCCC<br><u>AGTGAGCGTT</u> |
| BV42°C13 | <u>AACGCTCACT</u> CCXCC **AAGAAGAA** CCCCC **CCC** CCCCC **CCCC** CCCCC<br><u>AGTGAGCGTT</u> |
| BV42°C14 | <u>AACGCTCACT</u> CCCXC **AAGAAGAA** CCCCC **CCC** CCCCC **CCCC** CCCCC<br><u>AGTGAGCGTT</u> |
| BV42°C15 | <u>AACGCTCACT</u> CCCCX **AAGAAGAA** CCCCC **CCC** CCCCC **CCCC** CCCCC<br><u>AGTGAGCGTT</u> |
| BV42°C28 | <u>AACGCTCACT</u> CCCCC **AAGAAGAA** CCCCX **CCC** CCCCC **CCCC** CCCCC<br><u>AGTGAGCGTT</u> |
| BV42°C31 | <u>AACGCTCACT</u> CCCCC **AAGAAGAA** CCCCC **CCX** CCCCC **CCCC** CCCCC<br><u>AGTGAGCGTT</u> |
| BV42°C32 | <u>AACGCTCACT</u> CCCCC **AAGAAGAA** CCCCC **CCC** XCCCC **CCCC** CCCCC<br><u>AGTGAGCGTT</u> |

**Figure 2.**
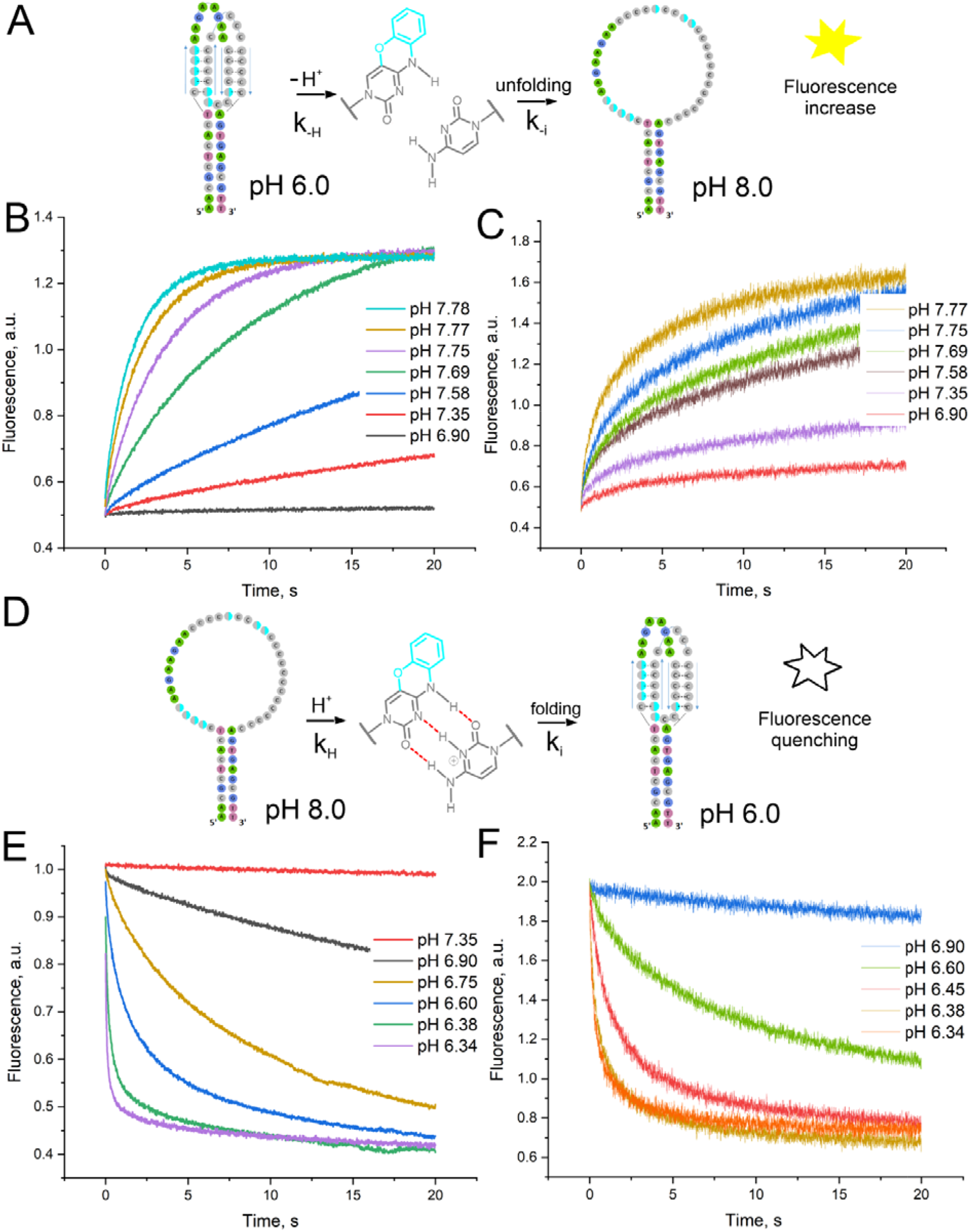
Kinetics of i-motif unfolding (A-C) and folding (D-F) of tC°-modified BV42 variants. Schemes of the processes (A and D). Curves for BV42°C14 (B, E) and BV42°C32 (C, F).

Stopped-flow kinetics were used to study i-motif folding and unfolding. The aptamers were assembled at pH 6.0 (a complete folding of the i-motif) or pH 8.0 (a complete unfolding of the i-motif part with retaining the duplex one [19]). Then a sample of the aptamer assembled at pH 6.0 (or 8.0) was mixed with the buffer at pH 8.0 (or 6.0) to initiate unfolding (or folding) of the i-motif module. Fluorescence of tC° is quenched due to the formation of a pair with C^+^ and stacking with neighboring C:C^+^ pairs that accompanied i-motif folding [20]. Examples of the kinetic curves are shown in Figure 2. i-Motif folding and unfolding are both multistep processes. The outer C:C^+^ pairs (see BV42°C15) get unpaired first during unfolding and paired at the end of i-motif folding. The C:C^+^ pairs inside the structure (e.g. BV42°C13) are more stable at neutral pH, being part of the i-motif core.

A simplified model was suggested to estimate the kinetics of i-motif folding and unfolding. Both processes can be described with the following scheme:

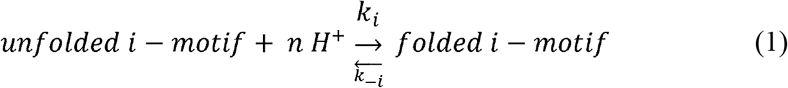

The fluorescence of tC° is quenched due to stacking and protonation [20]. So, one more intramolecular process is to be taken into account:

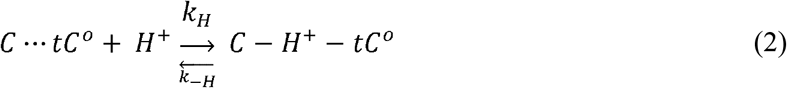

Data from stopped-flow kinetics experiments were fitted with the simplified model. The optimal number of protons (n) was 10 in full agreement with the predicted structure of the aptamer [19]. The derived parameters are provided in Tables 2 and 3. The values depend on the site of modification indicating local stability of i-motif. The folding and unfolding processes are not identical. The i-motif in the aptamer BV42 is surrounded by two loops from one side (the ‘loop’ side) and a duplex with one loop from another side (the ‘duplex’ side).

**Table 2.** Kinetic constants for i-motif unfolding. Kinetic constants of i-motif folding (k_i_) and unfolding (k_-i_), and derivative parameters (pH_i_) characterize the pH-stability of the i-motif near the tC° modification. Kinetic constants of C-H^+^-tC° pair formation (k_H_) and decay (k_-H_), and derivative parameters (pK_a_) characterize the inclusion of tC° modification into the i-motif structure.

| Code | $k_i, s^{-1}$ | $k_{-i}, s^{-1}$ | $pH_i$ | $k_H, s^{-1}$ | $k_{-H}, s^{-1}$ | $pK_a$ |
| --- | --- | --- | --- | --- | --- | --- |
| BV42°C12 | $(1.3 \pm 0.6) \cdot 10^{12}$ | $94 \pm 6$ | $7.01 \pm 0.03$ | $0.64 \pm 0.10$ | $0.041 \pm 0.018$ | $7.2 \pm 0.2$ |
| BV42°C13 | $(1.7 \pm 1.0) \cdot 10^{13}$ | $88 \pm 8$ | $7.13 \pm 0.10$ | $1.02 \pm 0.13$ | $0.030 \pm 0.003$ | $7.53 \pm 0.10$ |
| BV42°C14 | $(4.3 \pm 1.9) \cdot 10^7$ | $85 \pm 3$ | $6.57 \pm 0.03$ | $0.30 \pm 0.03$ | $0.017 \pm 0.08$ | $7.3 \pm 0.2$ |
| BV42°C15 | $(1.7 \pm 0.7) \cdot 10^8$ | $51 \pm 2$ | $6.65 \pm 0.02$ | $1.3 \pm 0.3$ | $0.037 \pm 0.012$ | $7.5 \pm 0.2$ |
| BV42°C28 | $(1.2 \pm 0.8) \cdot 10^{12}$ | $60 \pm 6$ | $7.03 \pm 0.05$ | $2.7 \pm 0.2$ | $0.19 \pm 0.04$ | $7.15 \pm 0.12$ |
| BV42°C31 | $(9 \pm 5) \cdot 10^5$ | $20.9 \pm 1.5$ | $6.46 \pm 0.10$ | $2.1 \pm 0.3$ | $0.248 \pm 0.011$ | $6.93 \pm 0.07$ |
| BV42°C32 | $(9 \pm 3) \cdot 10^{14}$ | $102 \pm 7$ | $7.29 \pm 0.03$ | $4.4 \pm 0.9$ | $0.41 \pm 0.04$ | $7.03 \pm 0.13$ |

**Table 3.** Kinetic constants for i-motif folding. Kinetic constants of i-motif folding (k_i_) and unfolding (k_-i_), and derivative parameters (pH_i_) characterize the pH-stability of the i-motif near the tC° modification. Kinetic constants of C-H^+^-tC° pair formation (k_H_) and decay (k_-H_), and derivative parameters (pK_a_) characterize the inclusion of tC° modification into the i-motif structure.

| Code | $k_i, \text{s}^{-1}$ | $k_{-i}, \text{s}^{-1}$ | $\text{pH}_i$ | $k_H, \text{s}^{-1}$ | $k_{-H}, \text{s}^{-1}$ | $\text{pK}_a$ |
| --- | --- | --- | --- | --- | --- | --- |
| BV42°C12 | 3600±700 | $(2.6\pm0.5)\cdot10^6$ | 5.71±0.02 | 5.0±0.3 | 0.17±0.07 | 7.5±0.2 |
| BV42°C13 | 9400±1300 | 0.971±0.012 | 6.40±0.02 | 2.4±1.0 | 1.94±0.05 | 6.1±0.2 |
| BV42°C14 | 3100±300 | 20±10 | 6.22±0.10 | 1.9±0.2 | 0.54±0.06 | 6.55±0.10 |
| BV42°C15 | 1800±300 | $(2.4\pm0.4)\cdot10^{10}$ | 5.29±0.02 | 1.33±0.18 | 1.0±0.2 | 6.12±0.14 |
| BV42°C28 | 4000±800 | $(1.1\pm0.7)\cdot10^6$ | 5.76±0.10 | 5.3±0.5 | 4.8±0.6 | 6.04±0.10 |
| BV42°C31 | 5000±900 | $(5.0\pm0.5)\cdot10^{10}$ | 5.30±0.02 | 3.5±0.5 | n.d. | n.d. |
| BV42°C32 | 2200±200 | $(1.5\pm0.3)\cdot10^8$ | 5.52±0.02 | 0.0163±0.013 | 0.028±0.002 | 5.77±0.07 |

The unfolding of the i-motif begins from the ‘loop’ side at pH>6.5, whereas the ‘duplex’ side is stable up to pH 7.0. The unfolding of the i-motif occurs with nearly the same rate, k_-i_∼100 s^−1^. The main differences are in the kinetic constants of i-motif folding. The rate constants for the ‘duplex’ side are in the range 10^12^-10^14^ s^−1^, whereas the rate constants for the ‘loop’ side are in the range 10^7^-10^8^ s^−1^. Thus, the accidentally distorted i-motif immediately restores if the distortion was near the ‘duplex’ side; and the structure restores 5 orders of magnitudes more slowly when the distortion was near the ‘loop’ side. Modification of the residue from the loop (BV42°C31) results in 10-100 lower rates of i-motif folding compared to the ‘loop’ side, indicating low stability of the outer C-H^+^-C pairs.

The folding process has another mechanism. Here, the kinetic constants for i-motif unfolding are indicative. The k_-i_ are minimal in the center of the nascent i-motif (BV42°C13 and BV42°C14), being in the range of 1-20 s^−1^. Next, the duplex part is assembled (k_-i_ ∼10^6^-10^8^ s^−1^). The outer C-H^+^-C pairs are formed at the last step of i-motif folding (BV42°C15 and BV42°C31).

Aptamer BV42 contains multiple base groups with pK_a_ in the 5.8-7.5 range. The upper limit is rather high compared with free cytosine pK_a_=4.7 [23]. However, pK_a_ for structured oligonucleotides can be much higher, e.g. pK_a_ of a GC pair was estimated to be of 7.2 [24].

### Ag^+^ is a poor competing ligand for the i-motif

Since unfolded i-motif of BV42 could bind Ag^+^ ions [19], a competition between H^+^ and Ag^+^ ions under i-motif-promoting and disrupting conditions was studied in detail. Stopped-flow kinetics were used to estimate the interaction of the aptamer with Ag^+^ ions. The interaction was tested at various pH levels in the range from 6.0 to 8.0. The major changes in tC^0^ fluorescence were observed at pH 8.0 and 7.3 (Figure 3), whereas pH 6.0, 6.5 and 7.0 caused minimal, if any, changes in fluorescence intensity (Supplementary Figure 10). The kinetic and equilibrium dissociation constants can be calculated only for several samples (Table 4). Two fluorescent derivatives of the BV42 aptamer were studied. BV42°C13 contained a fluorophore inside the i-motif core, whereas BV42°C32 had a fluorophore at the end of the i-motif. Notably, these two derivatives were not equal in Ag^+^ binding, which indicates competition between H^+^ and Ag^+^ ion binding with BV42. The folded i-motif has no affinity to Ag^+^ ions. The Ag^+^ ions are not able to displace H^+^ from the i-motif structure, as dissociation constants of C-Ag^+^-C are 2 order of magnitude higher compared to C-H^+^-C (Tables 2-4).

**Table 4.** Equilibrium dissociation constants (*K*_*D*_) and stoichiometry (*n*) of complexes of BV42 aptamer with Ag^+^ ions at various pH. Two derivatives were studied, namely, BV42°C32 (i-motif terminus) and BV42°C13 (i-motif core).

|  | 8.0 | 7.3 | 7.0 | 6.5 | 6 |
| --- | --- | --- | --- | --- | --- |
| i-motif terminus | $K_D=31\pm8\ \mu\text{M}$<br>$n=4.9\pm0.7$ | $K_D=59\pm8\ \mu\text{M}$<br>$n=1.7\pm0.2$ | $K_D=79\pm10\ \mu\text{M}$<br>$n=1.0\pm0.2$ | no affinity | no affinity |
| i-motif core | $K_D=34.9\pm1.4\ \mu\text{M}$<br>$n=8.4\pm1.4$ | no affinity | no affinity | no affinity | no affinity |

**Figure 3.**
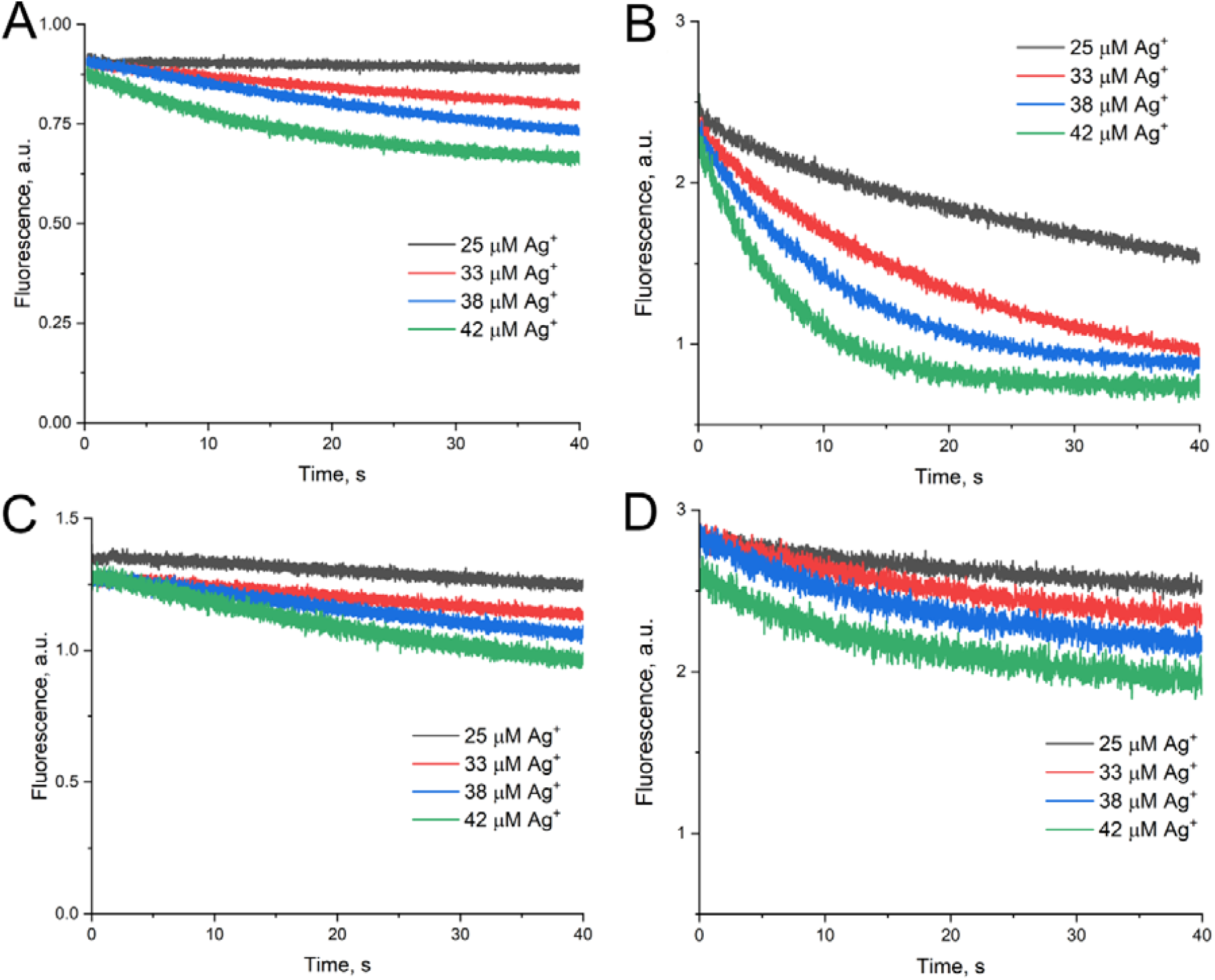
Kinetics of BV42°C13 (A, C) and BV42°C32 (B, D) aptamer complexation with Ag^+^ at pH 8.0 (A, B) and 7.3 (C, D).

The i-motif structure of BV42 has a stable core at pH 7.0-7.3, whereas the i-motif terminus is less stable (see previous subsection). Thus, the differences in dissociation constants of the BV42-Ag^+^ complex for two fluorescent derivatives are understandable. The two derivatives are rather similar at pH 8.0, when the i-motif is unfolded; both derivatives had K_D_=31-35 µM and the number of Ag^+^ ions (*n*) coordinated was 5-8. At pH 7.3, the i-motif core is assembled, providing no affinity to Ag^+^, whereas the i-motif terminus binds Ag^+^ with 2-times higher *K*_*D*_=59 µM and *n* of nearly 2. The affinity and binding site count decreased with decreasing of pH. At pH 7.0, *K*_*D*_=79 µM and *n* of nearly 1, further acidifying completely abolished the affinity to Ag^+^.

Isothermal titration calorimetry (ITC) was used to make a bulk estimation of the equivalence of Ag^+^ binding sites in the unfolded i-motif at pH 8.0 (Figure 4). The formation of complex resulted in an endothermal effect of nearly 4 kcal per 1 mole of Ag^+^ ions. The ITC data indicate several non-equal binding sites for Ag^+^ inside the BV42 structure (Figure 4A, C). The data were fitted with a model of 2 sets of sites that provided an averaged K_D_=15±7 µM with n=3.2±1.3. These data are rather close to calculations based on stopped-flow data (Table 4). Notably, the number of Ag^+^ binding sites is much lower than the maximum number of C-Ag^+^-C pairs. A more detailed study of multiple fluorescent derivatives can be used for accurate estimation of each Ag^+^ binding site (see the previous subsection). However, further study goes beyond the scope of this article.

**Figure 4.**
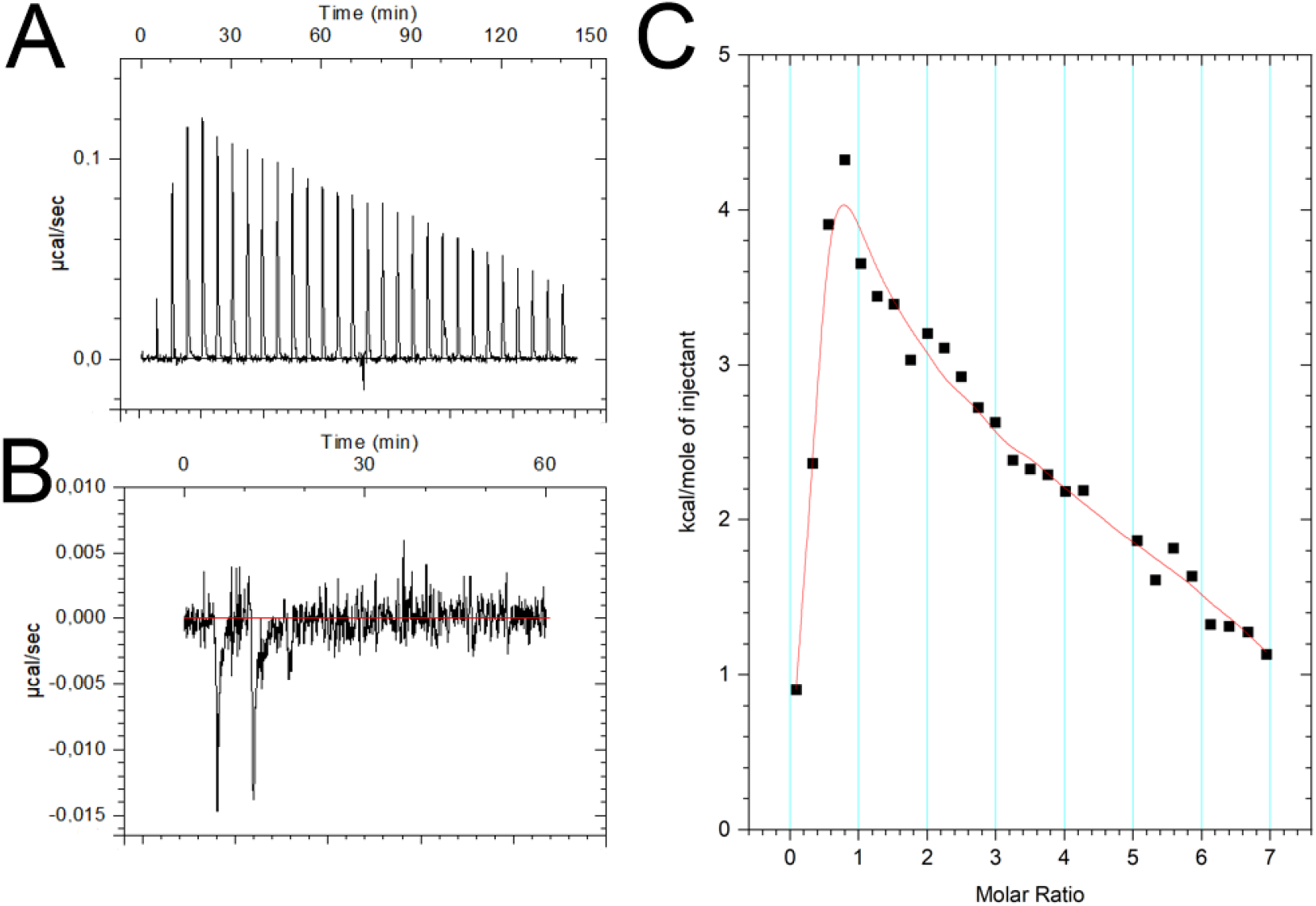
Isothermal titration calorimetry of Ag^+^ binding with BV42. Thermal effects at physiological ionic strength (A) and in water (B); normalized and fitted curve from subset (A) is shown in subset (C).

The thermodynamic parameters were also estimated. The molar enthalpy of the reaction of complex formation was estimated to be ΔH°_298_ =11±6 kcal/mol, and the molar entropy was ΔS °298 =60 cal/mol. The resulting change in free Gibbs energy for the reaction of complex formation was −7±3 kcal/mol at pH 8.0. The process is entropy-driven, which may be connected to the stacking of heterocycles with water being expelled from the hydrophobic interface.

Using ITC, an additional effect was revealed: the ability of Ag^+^ to bind the BV42 aptamer depends on the ionic strength of the buffer. The inability of the aptamer to bind Ag^+^ in water suggests that ionic strength (Figure 4B) might contribute to the stabilization of the duplex with C-Ag^+^-C pairs.

## Conclusions

Taken together, these data indicate that duplex context did not promote i-motif assembly. However, the assembled i-motif is more stable due to additional interactions with and spatial constrains imposed on the sugar-phosphate backbone by the duplex module. i-Motif unfolding was hindered near the duplex junction compared to the opposite ‘loop’ side. At the same time, the i-motif-duplex junction is not rigid allowing partial unfolding of the i-motif for another ligand binding (Ag^+^). The ligand binding is clearly pH-dependent; thus, the stability of the i-motif-duplex junction increases strongly at pH 6. These data reveal an additional variable in i-motif-dependent regulation. The i-motif could be completely assembled, partially folded with a large mobility in the i-motif-duplex junction, or completely unfolded.

## Materials and Methods

### Materials

Inorganic salts, acids, alkaline and tris were purchased from AppliChem GmbH (Darmstadt, Germany). AgNO_3_ was purchased from Sigma-Aldrich (St. Louis, MO, USA). Buffers with various pH were prepared based on conventional phosphate buffered saline (PBS, 10 mM Na_2_HPO_4_, 1.8 mM KH_2_PO_4_, 137 mM NaCl, 2.7 mM KCl pH 7.35) with addition of NaOH or HCl solutions. The target pH values were pH 6.0, 6.5, 7.0, 7.3 and 8.0. pH values were controlled with a ST20 pH-meter (Ohaus Corporation, Parsippany, New Jersey, United States). All solutions were prepared using ultrapure water produced by Millipore (Merck Millipore, Burlington, Massachusetts, United States).

### Oligonucleotides and sample preparation

Oligonucleotides were synthesized using commercially available reagents by a solid-phase phosphoramidite method, followed by purification with high performance liquid chromatography. The tC° phosphoramidite was prepared according to the reported procedure [25]. The sequences and sites of modification are provided in Supplementary Table 1. Typically, 2 or 5 µM solution of the aptamer in PBS buffer (pH 6.0, 6.5, 7.0, 7.3 or 8.0) was heated at 95°C for 5 minutes. The solutions were used after cooling to room temperature. The solutions were diluted with PBS buffer with the same pH, if necessary.

### Kinetic measurements and calculations

Stopped flow kinetics experiments were carried out using a MOS-200 rapid kinetics optical system equipped with an SFM-3000/S stopped-flow mixer (BioLogic Science Instruments, Seyssinet-Pariset, France). For i-motif folding, 5 µM aptamer solution in the buffer at pH 8.0 was placed in one syringe, whereas the buffer at pH 6.0 was placed in another syringe. For i-motif unfolding, 5 µM aptamer solution in the buffer at pH 6.0 was placed in one syringe, whereas the buffer at pH 8.0 was placed in another syringe. For coordination of Ag^+^ ions, 5 µM aptamer solution in the buffer at pH 6.0, 7.3 or 8.0 was placed in one syringe, whereas the same buffer with 50 µM AgNO_3_ was placed in another syringe. The solutions were mixed in different stoichiometries. Estimated dead times were in the range of 1.1-1.7 ms. The excitation wavelength was 370 nm; the emission spectrum was cut off with a 455 nm filter. The fluorescence was averaged for 2 ms (1 point) during 20-40 s. The curves are provided in Supplementary Materials (Figures S1 and S2).

The quenching of the fluorescence during i-motif folding was described with equations (1) and (2). The concentrations of folded (iM) and unfolded i-motif (uM) are connected with the total concentration of the oligonucleotide (iM_0_) through the fraction of folded i-motif:

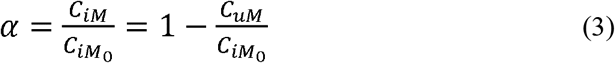

In general, the changes in fluorescence intensity can be represented as:

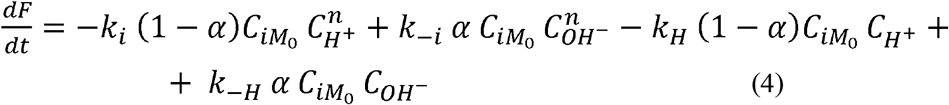

The concentration of hydroxyl ions was introduced to reflect the pH-dependent unfolding of the i-motif structure. Two derived parameters were used. The initial rate of the i-motif folding was derived from equation (4) given α→0 if t→0:

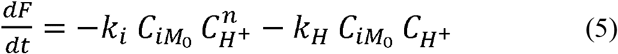

The equilibrium fraction of folded i-motif can be derived keeping dF/dt=0 and recalculating the concentration of hydroxyl ions in terms of the concentration of protons:

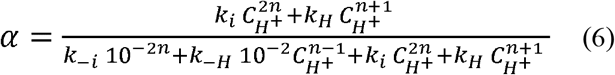

The initial rate of the i-motif unfolding was derived from equation (4) given α→1 if t→0:

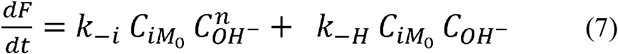

The equilibrium fraction of folded i-motif for unfolding process can be described with equation (6). Equations (5)-(7) were used to fit the kinetic curves acquired with stopped-flow kinetics. The first derivative was calculated for the first second of the reactions. This value was plotted versus concentration of protons and fitted with equations (5) and (7) for folding and unfolding, respectively. The equilibrium fraction of folded i-motif was calculated from exponential fit of the curves. The fluorescence at pH 6.0 or 8.0 for folded and unfolded i-motifs, respectively, were used as values for α=1 and α=0, respectively. The equilibrium fraction of folded i-motif was plotted versus concentration of protons and fitted with equation (6) for both folding and unfolding. The fitting curves are shown in Supplementary Materials (Figures S3-S). The fitted parameters are shown in Tables 2 and 3. The kinetic constants were recalculated in terms of the equilibrium constants in the following way:

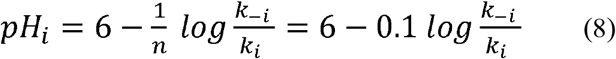

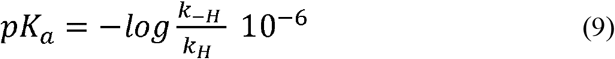

Ag^+^ binding was estimated using a simplified model. The fluorescence change can be described with the following function:

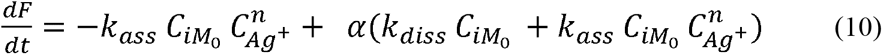

The association (*k*_*ass*_) and dissociation (*k*_*diss*_) rate constants can be estimated from the boundary conditions. The fraction of the complex (α) can be assumed to be 0 at the initial stages of the reaction:

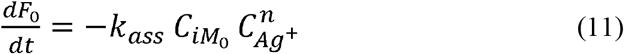

The fraction of the complex (α) can be assumed to be 1 at the final stages of the reaction:

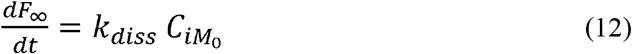

The stoichiometry and equilibrium dissociation constant can be estimated using the following equation:

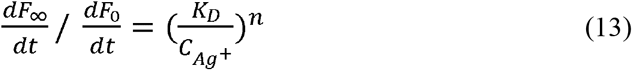

### Isothermal titration calorimetry

Tris-HNO_3_-based buffers with nitrate salts were used to exclude AgCl formation during the experiment. Namely, 20 mM Tris-HNO_3_ at pH 8.0 as well as 20 mM Tris-HNO_3_ at pH 8.0 with 140 MM NaNO_3_ were used as reaction buffers. The aptamer was additionally purified using centrifugation in 5x-excess of the buffer using Vivaspin 500 polyestersulfone membranes with 3 kDa cutoff (Sartorius Stedium Lab Ltd, Sperry Way, Stonehouse, UK). 3 µM aptamer solutions in the reaction buffer were heated at 95°C for 5 minutes. The solutions were used after cooling to room temperature. 100 µM AgNO_3_ in the reaction buffer was used as an injectant.

Heat production during the binding of Ag^+^ ions with the aptamer was measured at 25°C using a VP-ITC calorimeter (MicroCal Ltd, Piscataway, New Jersey, USA). Titration was performed in sequential injections of 10 µl, the time interval between injections was 5 minutes. The measured heat values were corrected to take into account the effects of dilution of the aptamer and injectant.

## Acknowledgements

The stopped flow experiments were performed using the equipment of the MSU Shared Research Equipment Center ‘Subdiffractional microscopy and spectroscopy’ and purchased by the MSU as part of the Equipment Renovation Program (National Project ‘Science’). Salary support from the Government project 121031300037-7 is acknowledged. The authors are grateful to A. Korshun for a fruitful discussion of the manuscript.

