## Supplementary materials for "Kinetics of i-motif folding within the duplex context"

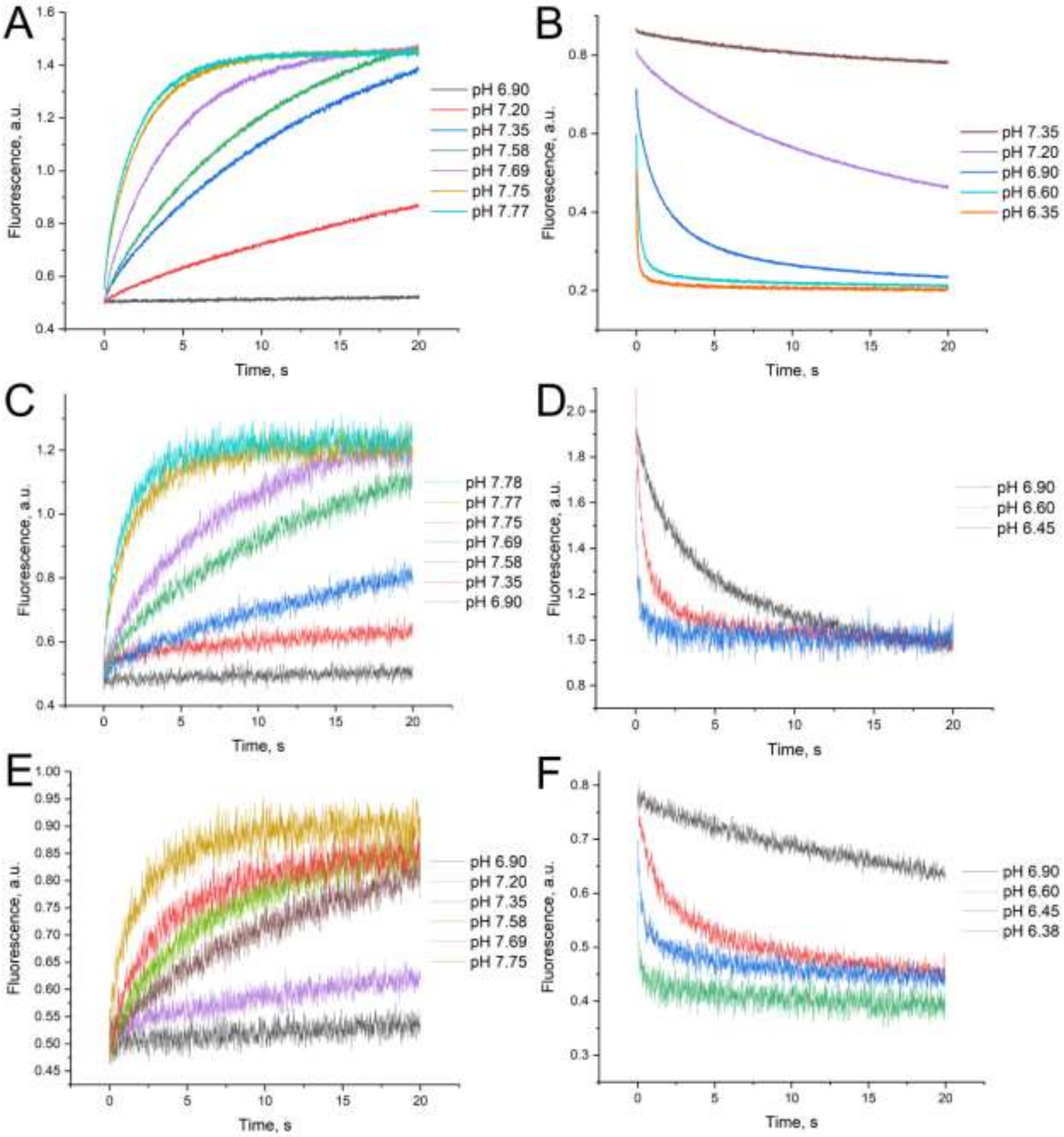


Figure S1. Kinetics of i-motif unfolding (A, C, E) and folding (B, D, F) of aptamers BV42°C12 (A, B), BV42°C13 (C, D) and BV42°C15 (E, F).

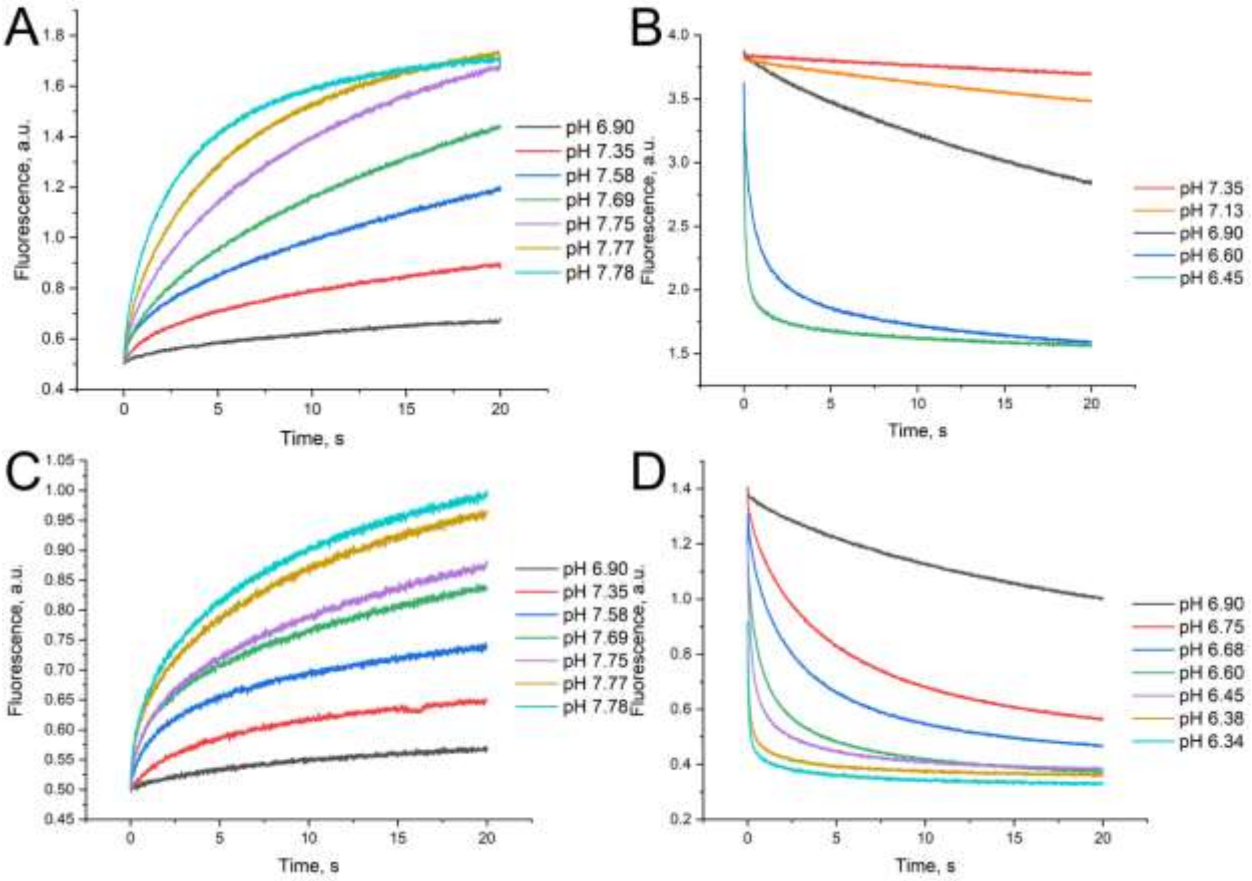


Figure S2. Kinetics of i-motif unfolding (A, C) and folding (B, D) of aptamers BV42°C28 (A, B) and BV42°C31 (C, D).

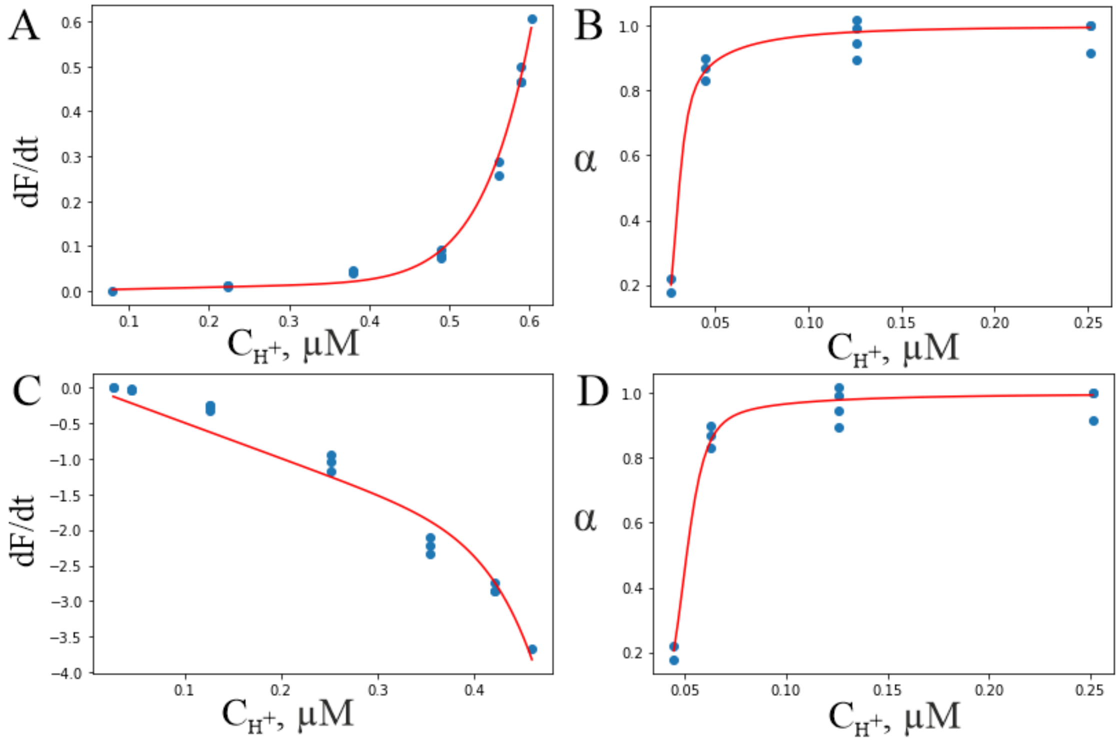


Figure S3. Fitting of kinetics of i-motif unfolding (A, B) and folding (C, D) of aptamer BV42°C12. Dependencies of the first derivative of fluorescence change over time (A, C) and equilibrium fraction of folded i-motif (B, D) on proton concentration.

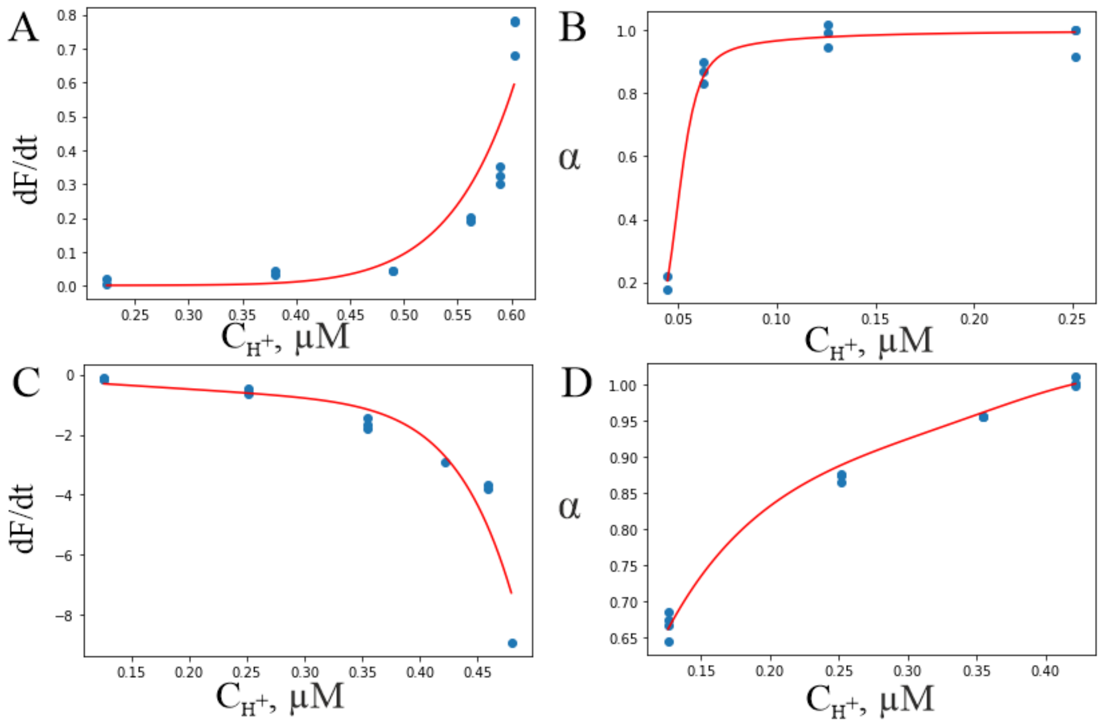


Figure S4. Fitting of kinetics of i-motif unfolding (A, B) and folding (C, D) of aptamer BV42°C13. Dependencies of the first derivative of fluorescence change over time (A, C) and equilibrium fraction of folded i-motif (B, D) on proton concentration.

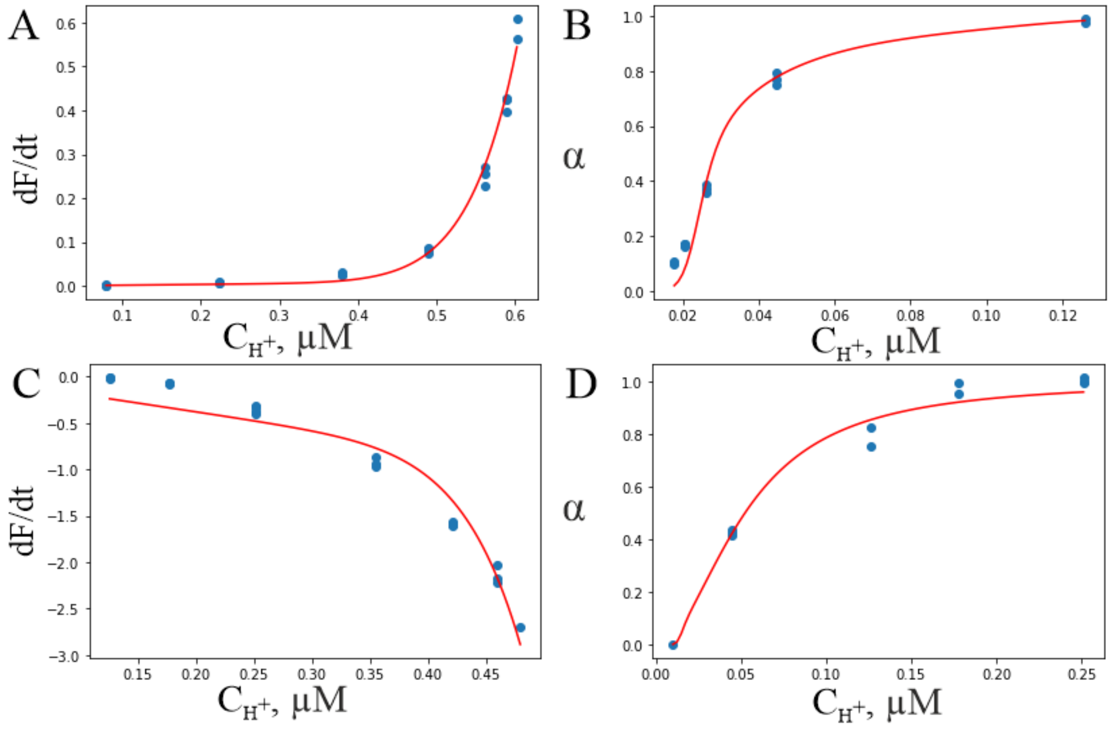


Figure S5. Fitting of kinetics of i-motif unfolding (A, B) and folding (C, D) of aptamer BV42°C14. Dependencies of the first derivative of fluorescence change over time (A, C) and equilibrium fraction of folded i-motif (B, D) on proton concentration.

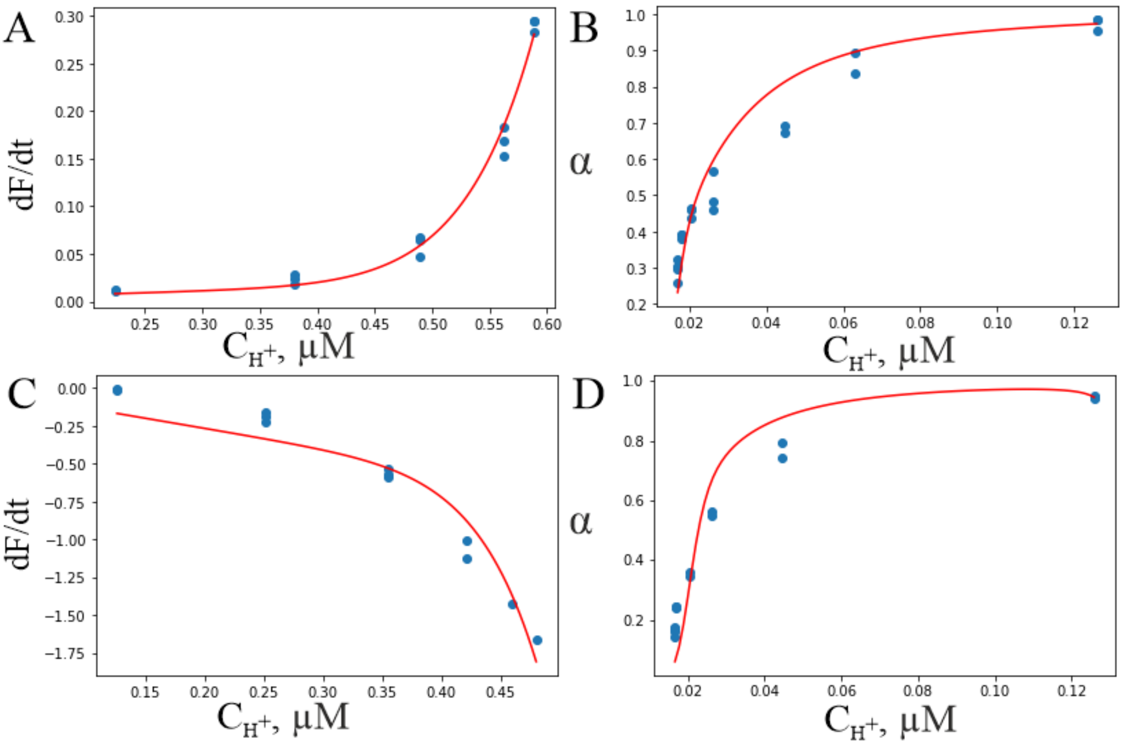


Figure S6. Fitting of kinetics of i-motif unfolding (A, B) and folding (C, D) of aptamer BV42°C15. Dependencies of the first derivative of fluorescence change over time (A, C) and equilibrium fraction of folded i-motif (B, D) on proton concentration.

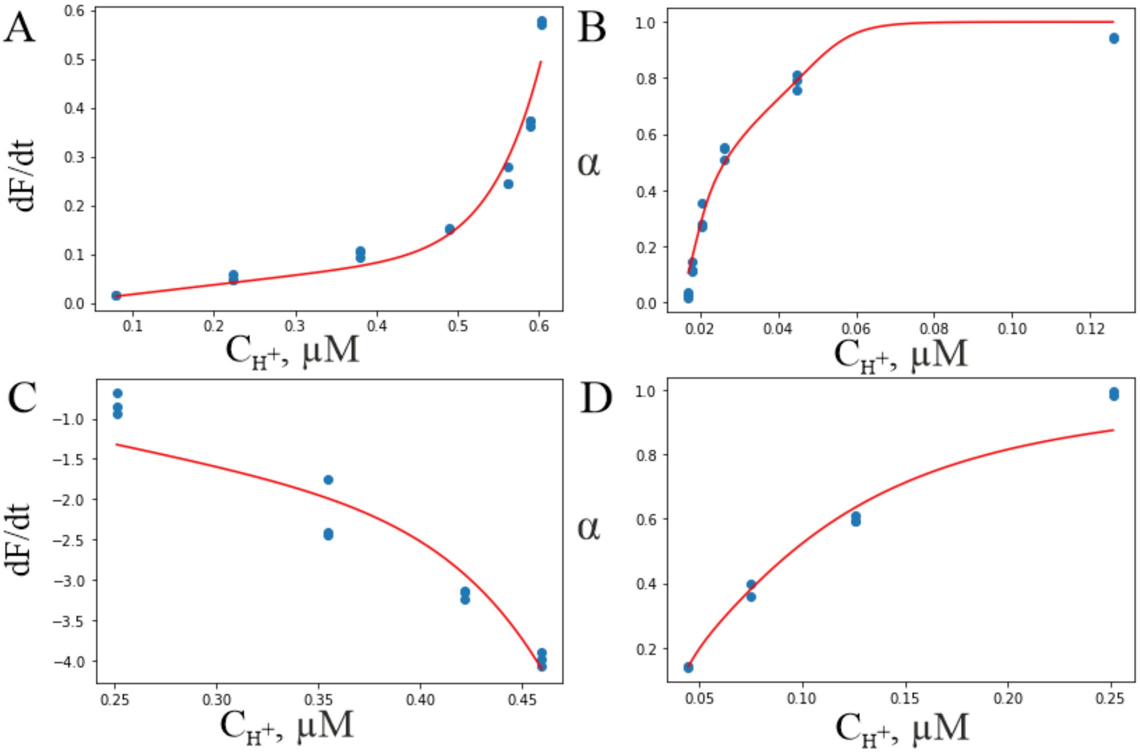


Figure S7. Fitting of kinetics of i-motif unfolding (A, B) and folding (C, D) of aptamer BV42°C28. Dependencies of the first derivative of fluorescence change over time (A, C) and equilibrium fraction of folded i-motif (B, D) on proton concentration.

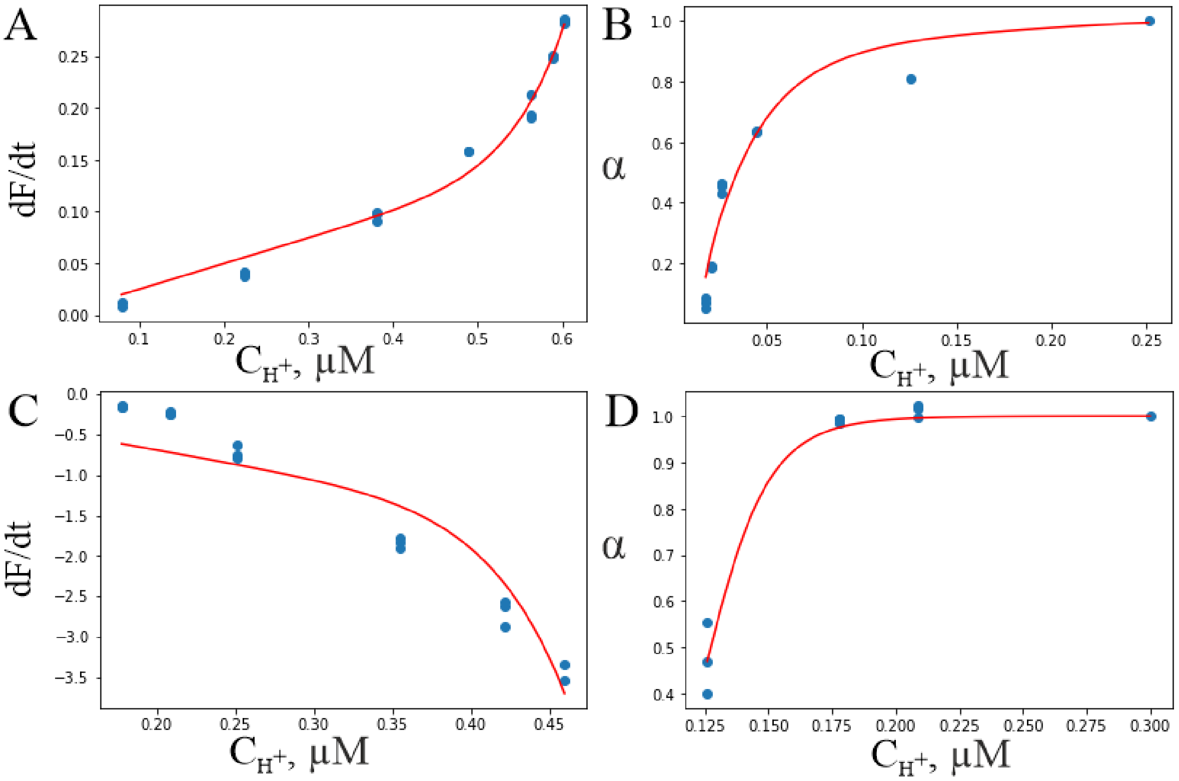


Figure S8. Fitting of kinetics of i-motif unfolding (A, B) and folding (C, D) of aptamer BV42°C31. Dependencies of the first derivative of fluorescence change over time (A, C) and equilibrium fraction of folded i-motif (B, D) on proton concentration.

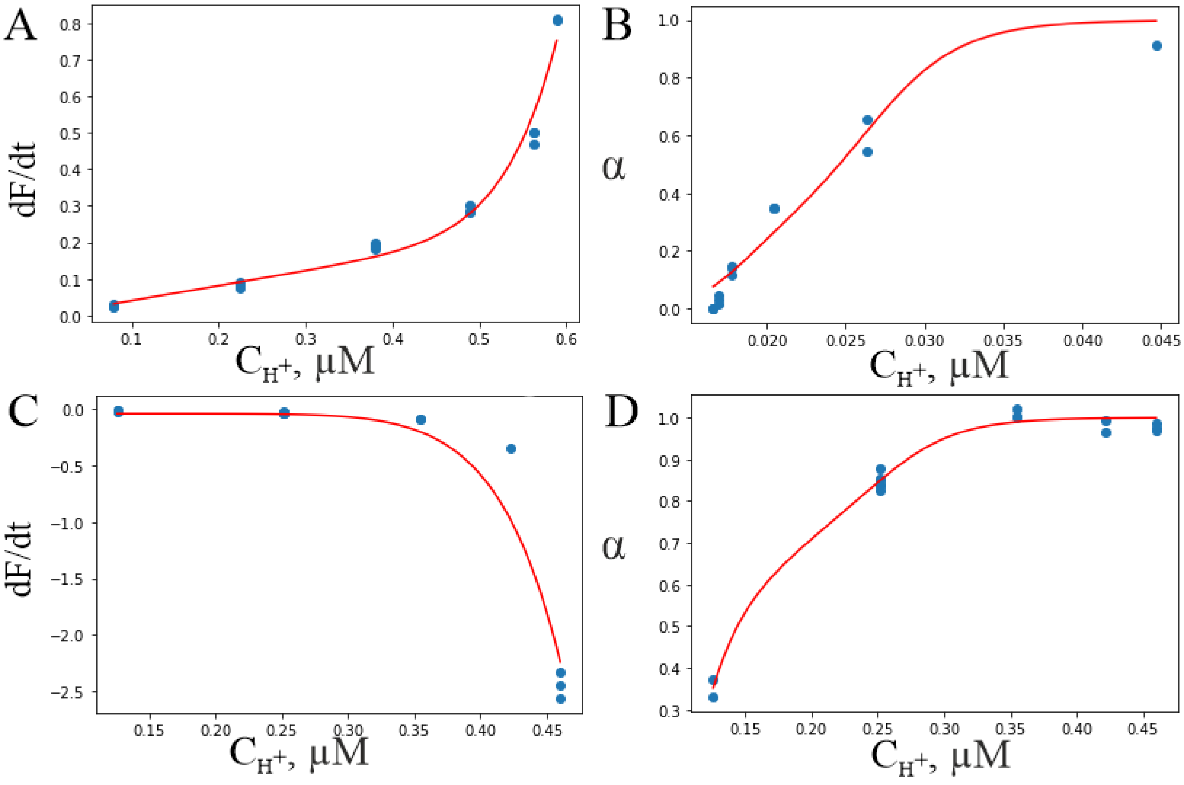


Figure S9. Fitting of kinetics of i-motif unfolding (A, B) and folding (C, D) of aptamer BV42°C32. Dependencies of the first derivative of fluorescence change over time (A, C) and equilibrium fraction of folded i-motif (B, D) on proton concentration.

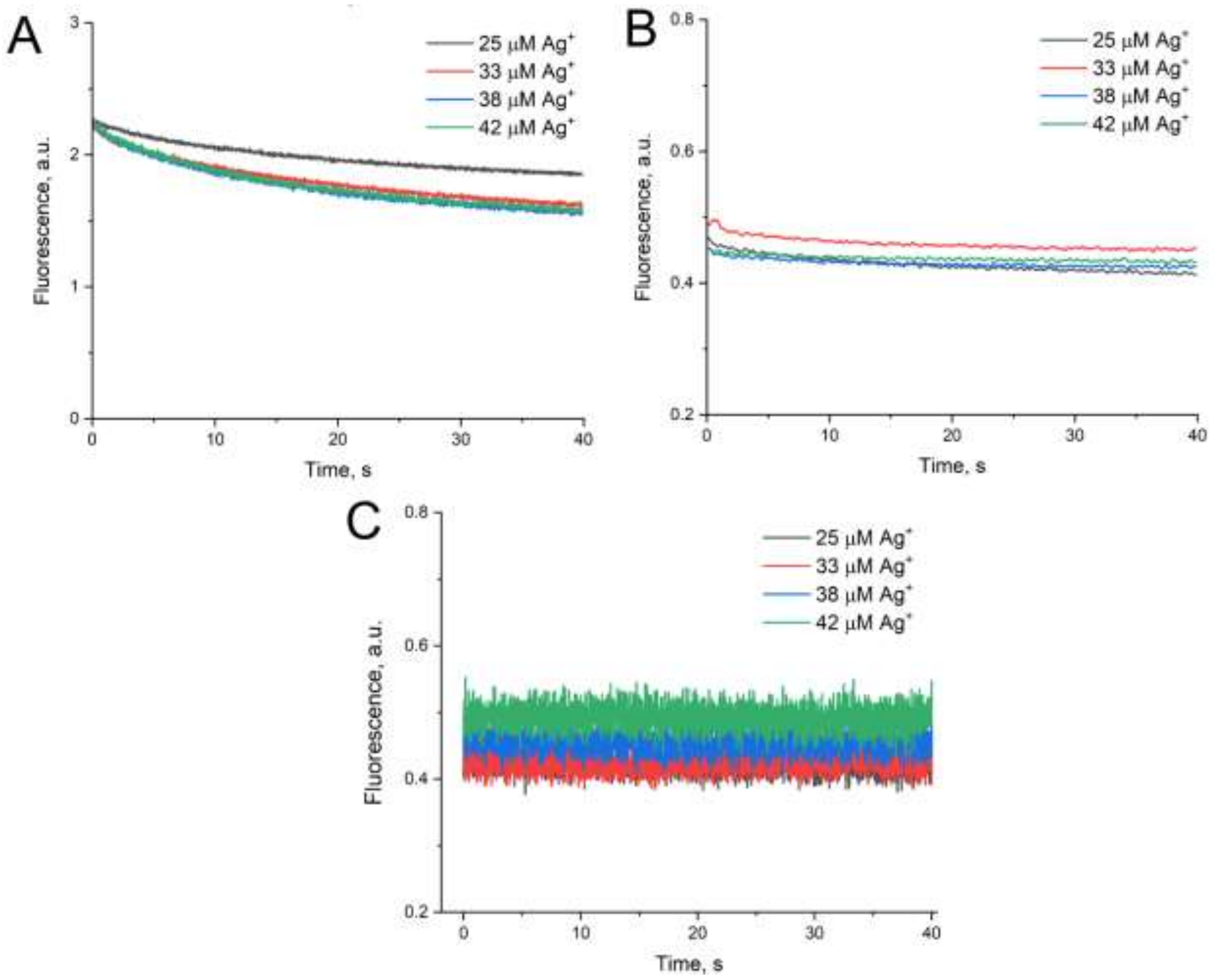


Figure S10. Kinetics of BV42°C32 aptamer complexation with $Ag^+$ at pH 7.0 (A), 6.5 (B) and 6.0 (C).
